# Mining ancient microbiomes using selective enrichment of damaged DNA molecules

**DOI:** 10.1101/397927

**Authors:** Clemens L. Weiß, Marie-Theres Gansauge, Ayinuer Aximu-Petri, Matthias Meyer, Hernán A. Burbano

**Affiliations:** Research Group for Ancient Genomics and Evolution, Department of Molecular Biology, Max Planck Institute for Developmental Biology, 72076 Tübingen, Germany; Department of Genetics, Stanford University, Stanford, CA, USA 94305; Department of Evolutionary Genetics, Max Planck Institute for Evolutionary Anthropology, 04103 Leipzig, Germany; Centre for Life’s Origins and Evolution, Department of Genetics, Evolution and Environment, University College London, WC1E 6BT, London, UK

**Keywords:** ancient DNA, metagenomics, authentication

## Abstract

**Background:** The identification of bona fide microbial taxa in microbiomes derived from ancient and historical samples is complicated by the unavoidable mixture between DNA from ante- and post-mortem microbial colonizers. One possibility to distinguish between these sources of microbial DNA is querying for the presence of age-associated degradation patterns typical of ancient DNA (aDNA). The presence of uracils, resulting from cytosine deamination, has been detected ubiquitously in aDNA retrieved from diverse sources, and used as an authentication criterion. Here, we employ a library preparation method that separates molecules that carry uracils from those that do not for a set of samples that includes Neandertal remains, herbarium specimens and archaeological plant remains.

**Results:** We show that sequencing DNA libraries enriched in molecules carrying uracils effectively amplifies age associated degradation patterns in microbial mixtures of ancient and historical origin. This facilitates the discovery of authentic ancient microbial taxa in cases where degradation patterns are difficult to detect due to large sequence divergence in microbial mixtures. Additionally, the relative enrichment of taxa in the uracil enriched fraction can help to identify bona fide ancient microbial taxa that could be missed using a more targeted approach.

**Conclusions:** Our experiments show, that in addition to its use in enriching authentic endogenous DNA of organisms of interest, the selective enrichment of damaged DNA molecules can be a valuable tool in the discovery of ancient microbial taxa.

## Background

DNA retrieved from historical or ancient samples is a complex mixture of molecules that contains not only endogenous host DNA, but also DNA from microorganisms that were present ante-mortem or that colonized the tissue post-mortem [1]. Therefore, all ancient DNA (aDNA) shotgun sequencing projects are metagenomic in nature. While earlier aDNA research has mostly focused on the evolution of animals and plants [2, 3], a growing number of studies are now centering on the identification and characterization of ancient pathogens and microbiomes [4]. Ancient microbes permit the replacement of indirect inferences about the past with direct observations of microbial genomes through time. By studying ancient pathogens, it has been possible to identify causal and/or associated agents of historical plant and animal disease outbreaks, as well as their spreading patterns throughout both space and time (e.g. [5, 6]). Beyond single taxa, another challenging endeavour is the characterization of shifts in composition of microbial communities over time. For example, dental calculus from hominids has been exploited as a source of ancient microbiomes and analyzed in the context of diet and lifestyle changes [7–9], whereas coprolites have been used to investigate ecological interactions between animals and microorganisms [10]. However, this approach is at its beginnings and the interactions of evolutionary processes in shaping microbiomes remain to be explored.

A major challenge for the study of aDNA in general, and ancient microbiomes in particular, is the presence of contaminating exogenous DNA, which makes distinction between *bona fide* ancient microbiome sequences and those of recent origin crucial. One of the most typical features of aDNA is the presence of uracils (Us) that originate from post-mortem deamination of cytosines (Cs), especially in single-stranded overhangs at molecule ends [11]. Uracils are read as thymines (Ts) by most DNA polymerases, which generates a characteristic increase in C-to-T substitutions at the end of aDNA sequences ([11], Figure 1D and 2A). This pattern makes aDNA damage distinguishable from biological sequence variation, and thus, its presence can be used as evidence for the authenticity of DNA sequences retrieved from ancient and historical material [12–14].

**Figure 1.**
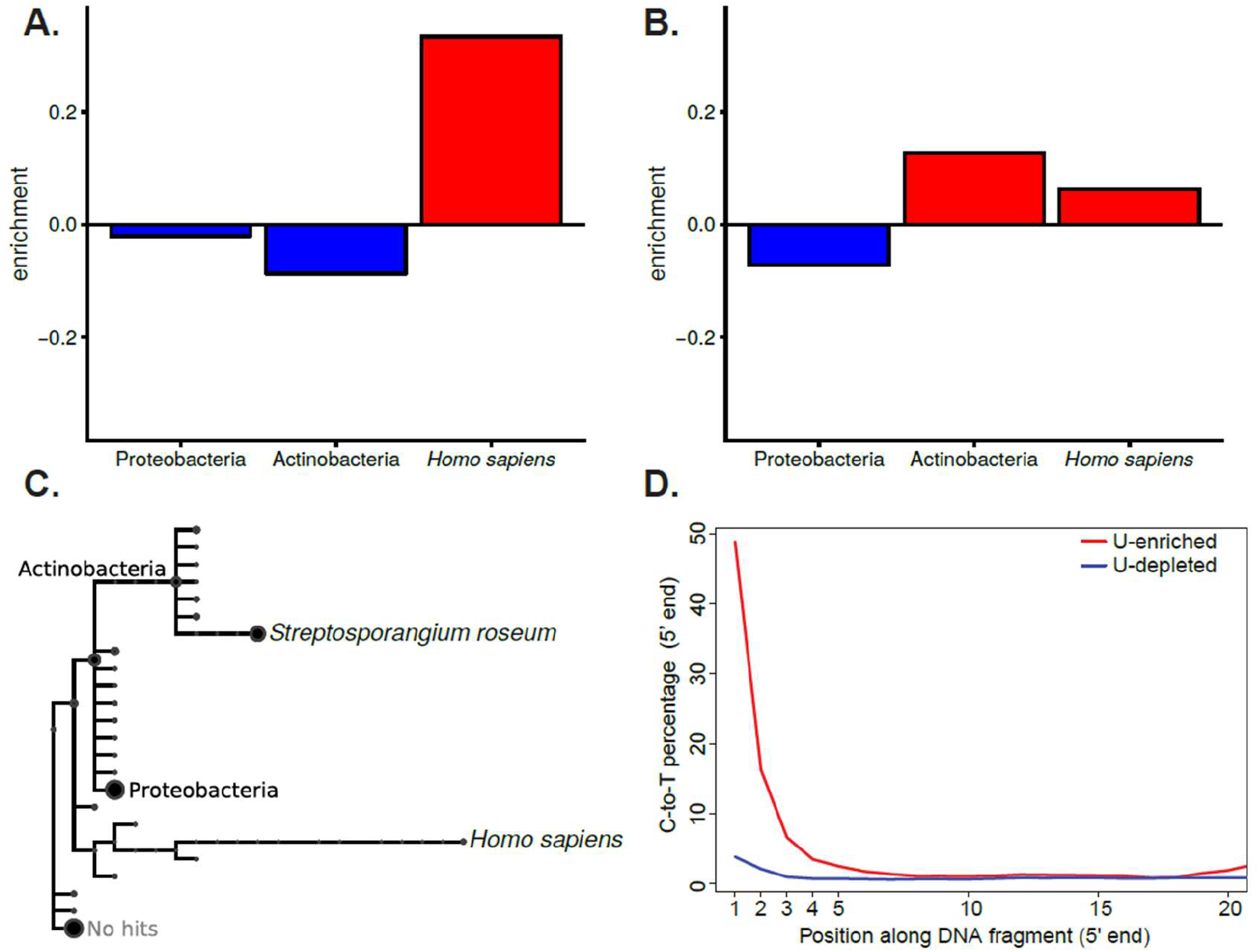
Relative enrichment, taxonomic assignment and substitution profiles of Neandertal-derived U-selected libraries. **A.** Relative enrichment (number of reads) in the U-enriched relative to the U-depleted fraction from Vindija Neandertal assigned to the phyla Actinobacteria and Proteobacteria, as well as to *Homo sapiens*. **B.** Relative enrichment (number of reads) in the U-enriched relative to the U-depleted fraction from Sidrón Neandertal assigned to the phyla Actinobacteria and Proteobacteria, as well as to *Homo sapiens*. **C.** Taxonomic tree of reads from Sidrón Neandertal assigned to different taxonomic levels. The size of the circle represents the amount of reads assigned to the node displayed in the tree or to any taxonomic level below it. Assignments to the phyla Actinobacteria and Proteobacteria, as well as the species *Streptosporangium roseum* and *Homo sapiens* are named in the taxonomic tree. Other taxonomic groups are either unlabeled or removed from the tree to increase clarity. **D.** Cytosine to Thymine substitutions at the 5’ end of reads aligned to S. *roseum* from the Sidrón Neandertal U-selected library (U-enriched and U-depleted fractions).

**Figure 2.**
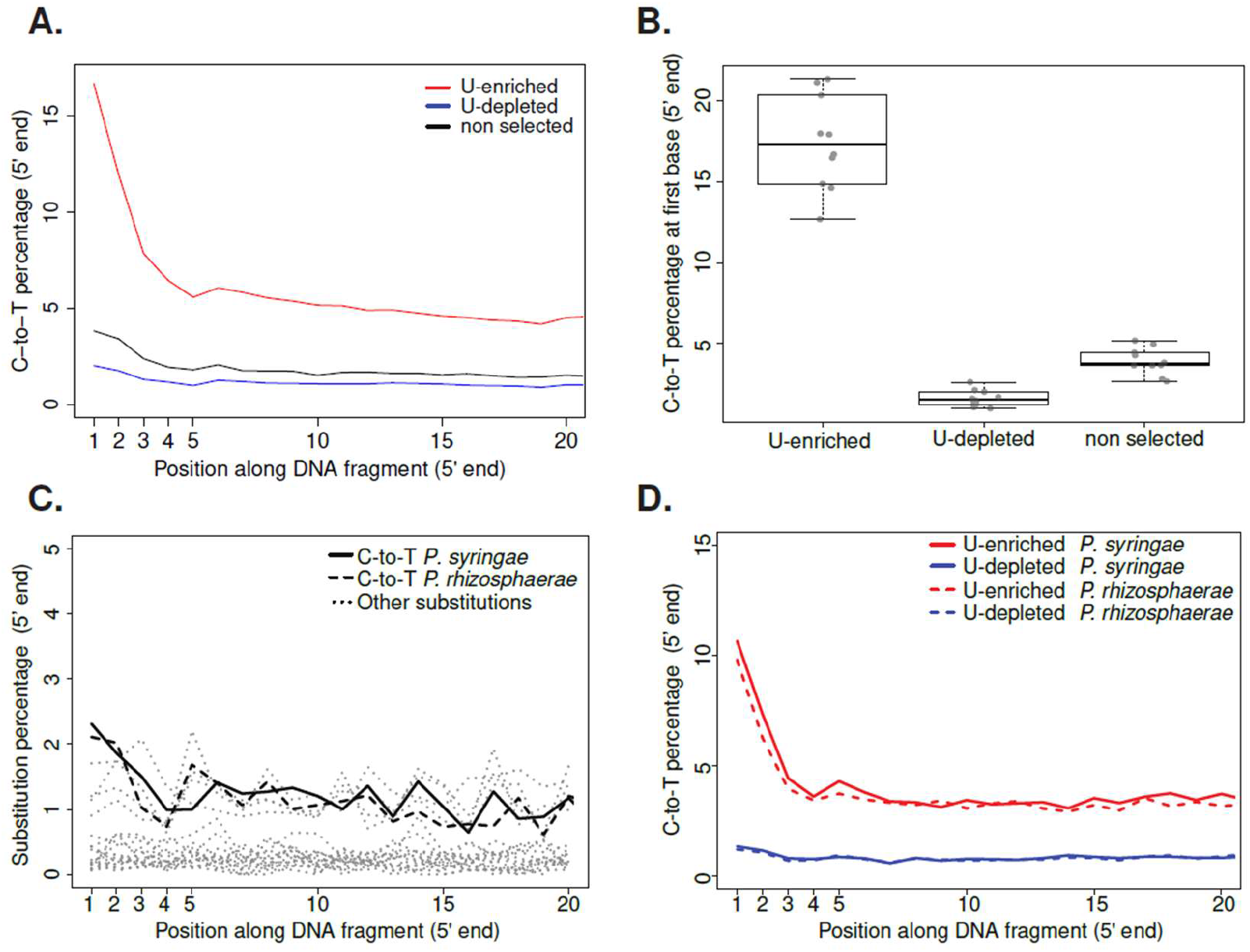
Patterns of cytosine to thymine (C-to-T) substitutions at the 5’ end of plant- and *Pseudomonas-derived* reads. **A.** C-to-T substitutions at the 5’ end of *Solanum tuberosum* sample KM177500 for a non-selected and U-selected library (U-enriched and U-depleted fractions). **B.** Distributions of C-to-T substitution percentage at first base (5’ end) for non-selected and U-selected libraries (U-enriched and U-depleted fractions). Median values are denoted as black lines and points show the original value for each individual sample. **C.** Substitution patterns at the 5’ end of *Pseudomonas syringae* and *Pseudomonas rhizosphaerae* mapped reads from a nonselected library from a *Solanum tuberosum* sample KM177500. **D.** Cytosine to Thymine substitutions at the 5’ end of *P. syringae* and *P. rhizosphaerae* mapped reads from a U-selected library (U-enriched and U-depleted fractions) from a *Solanum tuberosum* sample KM177500.

Recently, a single-stranded library preparation method (U-selection) was developed, which allows physical separation of uracil-containing molecules from non-deaminated ones [15]. In U-selection all library molecules are initially immobilized on streptavidin beads, to which molecules without uracils remain attached (U-depleted fraction), while uracil-containing molecules (originally deaminated) are released into solution (U-enriched fraction). U-selection was originally developed with the aim of increasing the amount of ancient hominid DNA (e.g. Neandertals) from a background of present-day human and microbial DNA [15]. However, the method seems to be specially suited to study microbiomes, due to the inherent difficulty to authenticate their ancient origin. This complication arises because microbes can colonize tissues at different times, resulting in different levels of deamination in ancient DNA molecules from different sources. Although sequences that carry terminal C-to-T substitutions can be selected *in silico* [16, 17], there are two factors that could hinder this approach. Firstly, low levels of deamination will reduce the number of molecules suitable for selection *in silico*. Secondly, high sequence divergence between samples and reference genomes can mask age-associated deamination signals thereby hinder authentication. Consequently, enriching for deaminated molecules during library preparation is fundamental to tackling these problems. As a proof-of-principle experiment, we used here U-selection in combination with taxonomic binning of Illumina sequenced reads to characterize the microbiomes of Neandertal bones (~39,000 years old), herbarium specimens (between 41 and 279 years old) and plant archaeological remains (~2,000 years old) (Table 1). Instead of exhaustively characterizing the composition of microbiomes in each sample, we aim at identifying authentic and highly abundant historical and ancient microbes through the presence of aDNA-associated damage patterns.

**Table 1.** Provenance of herbarium specimens and archaeological remains

| ID | Country of origin | Age | Species | Reference* |
| --- | --- | --- | --- | --- |
| KM177500 | UK | 171<sup>+</sup> | *Solanum tuberosum* | 1 |
| KM177497 | UK | 170<sup>+</sup> | *Solanum tuberosum* | 1 |
| BM000815937 | UK | 279<sup>+</sup> | *Solanum lycopersicum* | 2 |
| BH0000061459 | USA | 119<sup>+</sup> | *Arabidopsis thaliana* | 3 |
| OSU13900 | USA | 82<sup>+</sup> | *Arabidopsis thaliana* | 4 |
| NY1365364 | USA | 127<sup>+</sup> | *Arabidopsis thaliana* | 5 |
| NY1365375 | USA | 119<sup>+</sup> | *Arabidopsis thaliana* | 5 |
| CS5 | USA | 1852<sup>++</sup> | *Zea mays* | 6 |
| CS6 | USA | Undated | *Zea mays* | 6 |
| CS20 | USA | 1881<sup>++</sup> | *Zea mays* | 6 |
| El Sidrón 1253 | Spain | 39,000<sup>++</sup> | Neanderthal | 7 |
| Vindija 33.17 | Croatia | Undated | Neanderthal | 8 |
| Vindija 33.19 | Croatia | Undated | Neanderthal | 8 |

\*<sup>1</sup>Kew Royal Botanical Gardens; <sup>2</sup>Natural History Museum, London; <sup>3</sup>Cornell Bailey Hortorium; <sup>4</sup>Ohio State University Herbarium; <sup>5</sup>New York Botanical Garden; <sup>6</sup>Turkey Pen Shelter, UTAH, USA; <sup>7</sup>El Sidrón Cave, Spain; <sup>8</sup>Vindija Cave, Croatia.
<sup>+</sup>Calculated from collection dates (in years).
<sup>++</sup>B.P. (Before present years)

## Results

Our experiments were motivated by the previous observation that in some Neandertal samples, e.g. from El Sidrón, Spain, the proportion of Neandertal DNA fragments remains unchanged in both the U-depleted and U-enriched fractions, whereas in others, e.g. from Vindija Cave, Croatia, this proportion increased in the Uracil-enriched fraction [15]. It was hypothesized that the latter effect could have been due to differences in deamination, and hence in age, between Neandertal- and microbial-derived DNA fragments. To explore this effect further, we re-analyzed the previously generated Neanderthal sequence data from both sites by performing taxonomic binning of reads derived from the U-depleted and U-enriched fractions, instead of aligning them only to the human reference genome, as had been done previously. Reads aligning to the two most abundant bacterial phyla (Actinobacteria and Proteobacteria) from the Vindija Neandertals were enriched in the U-depleted fraction, while hominid reads were enriched in the U-enriched fraction (Figure 1A). This is in accordance with a previous study that reported absence of DNA damage in Actinobacteria derived from a Neandertal bone from Vindija cave [18]. In contrast, in reads obtained from the El Sidrón Neandertals, we found enrichment of both hominid and Actinobacteria reads in the U-enriched fraction, whereas Proteobacteria reads were enriched in the U-depleted fraction (Figure 1B). Overall, bacteria-derived reads were dominated by the Actinobacteria *Streptosporangium roseum* (Figure 1C), which showed almost 50% deamination at the first base in the U-enriched fraction (Figure 1D), suggesting its ancient origin. The analysis of reads derived from Neandertal bones illustrates how U-selection permits distinguishing between ancient bacteria enriched in the U-enriched fraction and more recent colonizers enriched in the U-depleted fraction.

In order to further evaluate the performance of U-selection in characterizing microbial communities in different sample types and with varying levels of deamination, we selected a set of plant samples including both herbarium specimens and archaeobotanical remains. In particular, we sought to investigate how the method performs in herbarium samples, which have much lower levels of deamination than for example Neandertal remains [19]. We extracted DNA from these plant samples and generated libraries using both a regular double-stranded (ds) approach [20], and U-selection [15]. Sequences from the dsDNA libraries, the most common library preparation method used nowadays, were then used as a baseline to evaluate depletion and enrichment of uracil-containing molecules (Figure 2A). U-selection successfully enriched for deaminated molecules in all plant samples, as shown by the much higher levels of deamination present in the U-enriched fraction compared with the dsDNA libraries and the U-depleted fraction (Figure 2A-B). The plant samples showed substantial variation in the content of endogenous DNA (2.8-91%), which was very similar between the U-depleted and U-enriched fractions, indicating similar levels of deamination between host- and microbe derived reads (Figure S1A). Assuming that plant- and microbial-derived DNA deaminate at a similar rate [19], this observation indicates that microbes found in our plant samples were present at the time of collection or colonized the tissue shortly thereafter, without substantial recent colonization. The percentage of reads (including host-derived reads) that could be taxonomically binned varied depending on the sample (Figure S1B) and, since the host genome was included in the nucleotide database, positively correlated with the percentage of host endogenous DNA (Figure S1E). The inability to taxonomically assign the vast majority of reads from samples with low endogenous DNA reflects the incompleteness of the reference database compared to the diversity of the microbiomes in those samples. Additionally, single stranded DNA library preparation methods as employed during Uracil enrichment generate shorter reads [21, 22], which are more difficult to map to a reference genome and to assign taxonomically to a nucleotide database. This is reflected in the higher percentage of reads mapped and assigned from the dsDNA library compared with shorter reads derived from both the U-depleted and U-enriched fraction (Figure S2A-B). Originally, it was reported that the U-enriched fraction shows a mild increase in GC-content [15], however in the plant libraries analyzed here we did not find a significant difference in GC-content between the U-depleted and U-enriched fractions (Figure S2C). In theory, since Us originate from Cs, the U-enriched fraction would be enriched for GC-rich species and GC-rich genomic regions within a given genome. However, as the enrichment would depend on the diversity of taxa present and their relative age difference, and hence difference in deamination, GC-biases, if any, are expected to be highly sample-dependent.

Given the low compositional complexity of microorganisms in the samples included in our proof-of-principle experiment, instead of centering our analyses on the compositional assessment of microbial communities, we investigated in detail samples in which a specific microbe or group of microbes were more prevalent based on read abundance.

Identifying highly abundant microbes, especially pathogens, is currently the most common use of ancient microbiomes [4]. We identified a large number of reads that were assigned to the bacterium *Pantoea vagans* in a potato (*Solanum tuberosum*) and a maize (Zea mays) sample (Figure 3A). In both samples we found patterns of C-to-T substitutions that suggest the historical nature of the sequenced reads (Figure 3B). Since *P. vagans* is a plant epiphyte [23], it is not entirely surprising to find it in two different plant species. We compared the potato and maize *P. vagans* with publicly available genomes using single nucleotide polymorphisms (SNPs) ascertained in these modern samples, and genotyped in the U-depleted fraction of either specimen. Our analysis linked the two historical strains to a distinct cluster of modern strains based on genetic similarity (Figure 3C). Based on a set of 432,891 SNPs, the two historical isolates showed 95% SNP identity between them, and an average of 92% SNP identity between historical and modern strains of the same cluster. Conversely, comparisons between historical strains and any modern strain of a different cluster showed only an average of 59% identity at variable positions.

In a potato sample where the pathogenic oomycete *Phytophthora infestans* was previously identified [6], we found a large portion of reads assigned to the bacterial genus *Pseudomonas*. While most reads were classified only to the genus level, some reads were assigned to either the species *Pseudomonas syringae* or *Pseudomonas rhizosphaerae* in different proportions (Figure S3). The limited length of aDNA fragments made it challenging to gain higher resolution of the taxonomic diversity within the *Pseudomonas* genus. Therefore, we performed de novo assembly using reads assigned to the genus *Pseudomonas*, aiming to generate longer sequences (contigs) that allow more detailed taxonomic classification. We aligned the contigs to the reference genomes of *P. syringae* and *P. rhizosphaerae*, which resulted in covering about 80% of both reference genomes (Figure S4). To reliably identify contigs that originate from either *P. syringae* or *P. rhizosphaerae*, we subsequently filtered for contigs that aligned uniquely to either reference genome (Figure S5A). Among these species-specific contigs, we found different k-mer coverage distributions in contigs aligning uniquely to either genome (Figure S5), an observation that reinforced our confidence in the presence of the two *Pseudomonas* species in this sample. Due to the high level of sequence divergence between the *Pseudomonas* in our sample and the reference genomes present in the database, it is difficult to assess typical deamination patterns in the dsDNA library (Figure 2C). However, we were able to examine damage patterns in both *Pseudomonas* species using the U-enriched fraction (Figure 2D), since the C-to-T signal is amplified and thus much higher than the basal level of substitutions.

**Figure 3.**
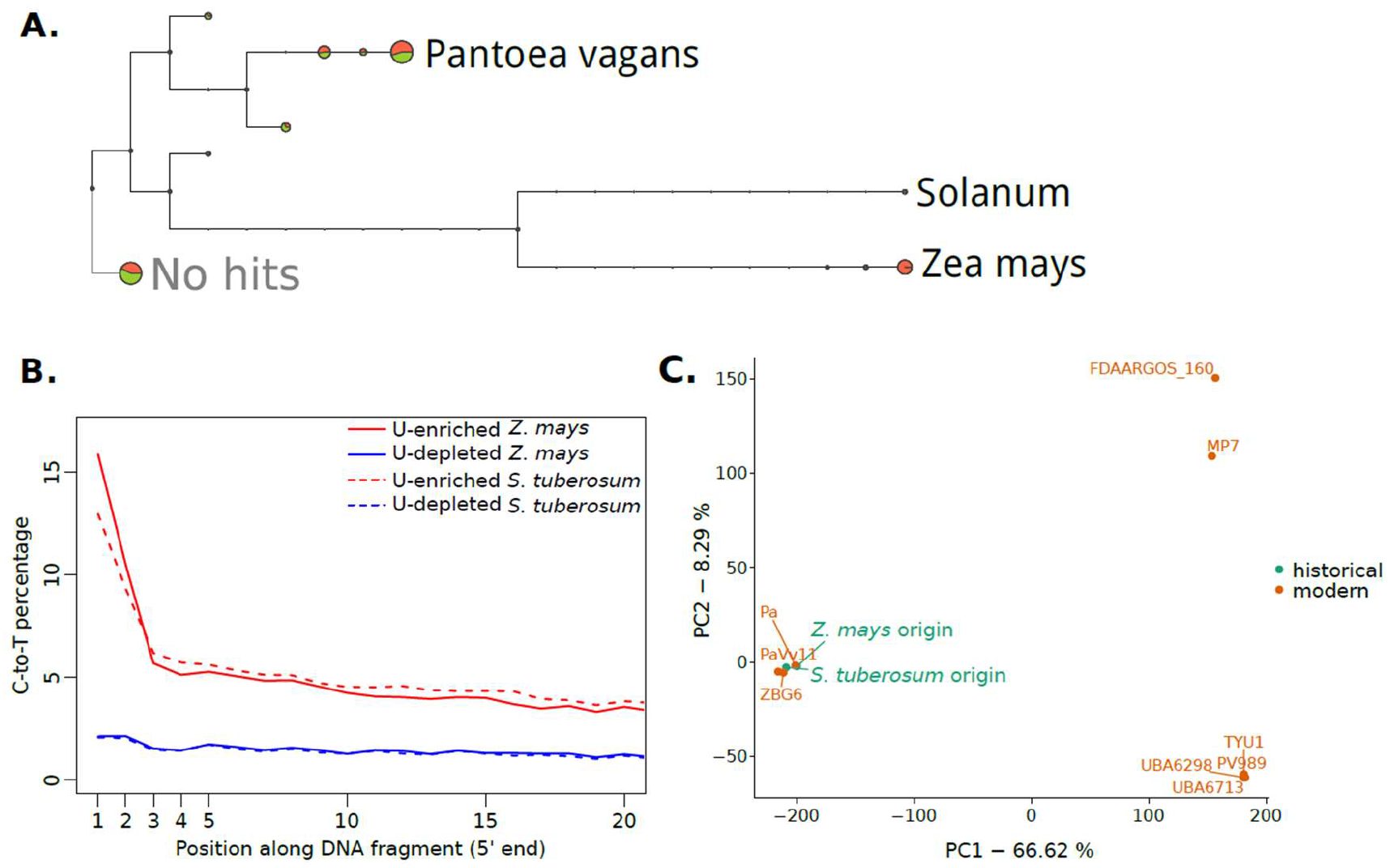
Characterization of the bacterium *Pantoea vagans* identified in *Zea mays* and *Solanum tuberosum* samples. **A.** Taxonomic tree of reads from *Solanum tuberosum* and *Zea mays* assigned to different taxonomic levels. The size of the circle represents the amount of reads assigned to the node displayed in the tree or to any taxonomic level below it. *S. tuberosum*- and *Z. mays*-derived reads are shown in green and orange, respectively. **B.** Cytosine to Thymine substitutions at the 5’ end of *P. vagans* for U-selected libraries (U-enriched and U-depleted fractions) from *Z. mays* and S. *tuberosum*. **C.** Principal component analysis of *P. vagans* from *Z. mays* and S. *tuberosum* samples, as well as nine publicly available genomes, based on single nucleotide polymorphisms. Numbers in axis labels indicate the percentage of the variance explained by each principal component (PC).

In summary, we showed here that the U-selection method selectively enriches for authentic microbial aDNA molecules in samples from plant and animal tissues with a wide-distribution of ages and deamination levels.

## Discussion

The utility of DNA from ancient and historical specimens is being recognized in a growing number of fields, ranging from human, animal and plant genetics, to microbiology and epidemiology of infectious diseases. Providing positive evidence for the authenticity of such ancient DNA from diverse sources is instrumental for all studies that make use of this resource, as mistaking modern contaminants or colonizers for ancient microbes can drastically influence the interpretation of results. Authentication is especially challenging when studying ancient microbes, due to their high genetic diversity and the incompleteness of reference databases. The method we employ and characterize here aids this process through the selective enrichment of molecules that carry signatures of age-associated degradation. For instance, in *P. vagans*, U-selection increases the fraction of molecules carrying a terminal C-to-T substitution at the 5’-end 2-3 fold over the library without enrichment, relative to the total number of molecules sequenced. We think that the application of U-selection for ancient microbiome research will be particularly useful in samples with minute levels of deamination, where the nucleotide divergence between samples and reference genomes will obscure the identification of the C-to-T pattern typical of aDNA, as well as in moderately or heavily deaminated samples which carry modern contaminants, since in those samples ancient taxa would be efficiently enriched. Since it is extremely difficult to differentiate between ante-mortem and early post-mortem colonizers based only on deamination patterns, it is fundamental to also evaluate the biological relevance of detected taxa by comparing them with reference modern microbiomes.

## Conclusions

We have applied the selective enrichment of uracil carrying DNA fragments to ancient DNA metagenomics. The presence of uracil is associated with age-associated degradation, and is commonly used as an authentication criterion when identifying ancient DNA sequences. We show that by assessing the relative abundance of microbial taxa in both the uracil enriched fraction and the uracil depleted fraction, it is possible to identify the presence of authentic ancient microbial taxa. Additionally, the “magnifying glass” effect of uracil enrichment on the detection of deamination patterns can help authenticating very divergent microbial taxa with moderate levels of degradation. Since there is no intersection of molecules between the U-selected and U-enriched fractions, splitting the sequencing efforts between the two fractions comes at no extra sequencing cost. All in all, we think that selective uracil enrichment is a valuable addition to ancient DNA metagenomics, both for de-novo taxon discovery as well as for cases where authenticity might be contentious.

## Methods

### Sequencing libraries from Neanderthal remains

We used DNA sequencing libraries from Neanderthal specimens (Table 1) prepared by [15], which were sequenced deeper for this study.

### DNA extracts from historical plant samples

Previously published DNA extracts from four different plant species were used for this study. These extracts were derived either from herbarium specimens (*Arabidopsis thaliana* [24], *Solanum tuberosum* [6], *Solanum lycopersicum* [6]), or from archaeobotanical remains (Zea *mays* [25]). Ages ranged from 41 to 279 years for herbarium samples, and 1852 to 1881 years for *Zea mays* samples (Table 1). The sequencing data for these samples is available on the European Nucleotide Archive under study number PRJEB30666.

### Sequencing libraries from historical plant samples

Three sequencing libraries were produced for each plant derived DNA extract, one using a double-stranded library preparation [20, 26] without enzymatic removal of uracils [27], and one for each fraction resulting from the single-stranded U-selection protocol [15].

### Sequencing and initial data processing

Since the length of aDNA molecules is often shorter than the read length of the sequencing platform, it is possible that a fragment of the aDNA molecule is sequenced by both the forward and reverse read, and also that a part of the adapter is sequenced [28]. Therefore, it is recommended to merge sequences based on the overlapping fraction sequenced by both forward and reverse reads [28]. We remove adapters and merged sequences using the software leeHom with the “--ancientdna” option [29] 91-99% of read pairs were merged (mean: 96.9%). Using the U-selection protocol, 81-97% of read pairs were merged (mean: 93.4%). Putative chimeric sequences were flagged as failing quality.

### Mapping of sequenced reads to their host genome

Merged reads were mapped as single-ended reads to their respective or most closely relative genome: *Zea mays* [30], *Arabidopsis thaliana* [31,32], *Solanum tuberosum* [33], *Solanum lycopersicum* [34], *Homo sapiens* (Genome Reference Consortium Human Build 37). The mapping was performed using BWA-MEM (version 0.7.10) with default parameters, which includes a minimum length cutoff of 30 bp [35].

### Metagenomics assignment of sequenced reads

Reads were aligned to the full non-redundant NCBI nucleotide collection (nt) database (downloaded January 2015) using MALT (version 0.0.12, [36]) in BlastN mode. The resulting RMA files were analyzed using MEGAN (version 5.11.3, [37]). The reads were assigned to the NCBI taxonomy using a lowest common ancestor algorithm [37].

### Mapping of sequenced reads to microbial genomes

Libraries were mapped to microbial reference genomes of interest, after the presence of certain taxa was detected during metagenomic assignment. Specifically, the references of *Streptosporangium roseum* [38], *Pseudomonas syringae* pv. *syringae* B728a [39], *Pseudomonas rhizosphaerae* [40] and *Pantoea vagans* [23] were used. Since mapping metagenomic libraries to bacterial reference genomes is very prone to false alignments, we used a different mapping strategy for these genomes. The mappings were performed with bowtie2 (version 2.2.4, [41]), with the settings “--score-min ‘L,-0.3,-0.3’ --sensitive-end-to-end” to increase stringency.

### Assessment of nucleotide substitution patterns

All types of nucleotide substitutions relative to the reference genome were calculated per library using mapDamage 2.0 (v. 2.0.2-12, [42]). The percentages of C-to-T substitutions at the 5’ end were extracted from the output file 5pCtoT_freq.txt produced by mapDamage.

### *Pantoea vagans* genomic variation

In order to reduce the effect of aDNA-associated C-to-T substitutions on variant discovery, we used exclusively the U-depleted fraction of libraries where *P. vagans* was detected in the metagenomic screening. The libraries were mapped to the *P. vagans* reference genome using BWA-MEM, to reduce reference bias and increase SNP discovery. False alignments from the metagenomic libraries posed a lesser problem here, as variants were ascertained based on modern material. Variants for historical samples were called for both libraries together using the bcftools (version 1.8, [43]) utilities mpileup (“bcftools mpileup -q 1 -I -Ou -f $REF $IN1 $IN2”) and call (“bcftools call --ploidy 1 -m -O -z”). Additionally, 11 assemblies of different contiguity were downloaded from NCBI (https://www.ncbi.nlm.nih.gov/genome/genomes/2707). These assemblies were aligned to the reference genome using minimap2 (version 2.10-r764, [44]) and its “asm20” parameter preset. Only strains with at least 80% reference coverage were kept for subsequent analysis (9/11, average reference coverage: 91%). The paftools utility, which is distributed with minimap2, was used to call variants from these alignments, with the parameter set “-l 2000-L 5000”. All resulting VCF files from modern samples were merged using bcftools’ merge utility with the parameter “--missing-to-ref”, assuming that those positions not called by paftools in any one sample were indeed reference calls. The merged VCF from modern material was then merged with the VCF from the two historical samples using bcftools (version 1.8, [43]), and filtered to include only full information, biallelic SNPs. This approach discovers sites, which are segregating in modern material, and have read data (be it reference, alternative or segregating sites) in both historical samples. The resulting VCF file was loaded into R using vcfR (version 1.7.0, [45]), and a PCA was produced by converting the information into a genlight object using adegenet (version 2.0.1, [46]) in R (version 3.3.3, [47]).

### *Pseudomonas* spp. assembly and evaluation

To evaluate the presence of *Pseudomonas* spp. strains in a *Solanum tuberosum* historic herbarium sample, we extracted from this library all reads that were taxonomically assigned to the *Pseudomonas* genus or to inferior taxonomic levels within it. These reads were then assembled using SPAdes (version 3.5.0) with default parameters [48]. The resulting contigs were filtered for a minimum length of 2Kb, which yielded 3,314 contigs with a total length of 16Mb (avg. contig: 4.8kb; N50: 5.3kb). We used the lastz (version 1.03.66, [49]) and Circos (version 0.64, [50]) interface of AliTV [51] to align these contigs to either the *P. syringae* or *P. rhizosphaerae* reference genome. We were able to align 72% of contigs to either one or both of these reference genomes in alignments of at least 1Kb. We then extracted all contigs which had alignments of at least 10Kb in length and were unique to one of the reference genomes. These contigs were again aligned to their corresponding reference using AliTV as described above. Using the “links” file produced by AliTV, we visualized all alignments with circlize [52]. Additionally, we used uniquely aligning contigs to assess their average kmer coverage during the assembly as reported by SPAdes.

## List of Abbreviations

aDNA: ancient DNA
VCF: Variant Call Format
PCA: Principal Component Analysis
dsDNA: double stranded DNA libraries
ssDNA: single stranded DNA libraries

## Declarations

## Ethics approval and consent to participate

Not applicable.

## Consent for publication

Not applicable.

## Availability of data and material

The sequencing data generated for this study is available on the European Nucleotide Archive under study number PRJEB30666.

## Authors’ contributions

CLW, MM and HAB conceived and designed the study. M-TG and AA-P generated sequencing data. CLW analyzed the data. CLW and HAB wrote the manuscript with input from all authors. All authors read and approved the final manuscript.

## Acknowledgements

We thank Verena Schuenemann for help in the laboratory; Patricia Lang, Claudia S. Burbano, members of the Research Group for Ancient Genomics and Evolution, and specially Talia Karasov for input on data analysis and comments on the manuscript; Janet Kelso for help with processing the raw sequencing data; Ivan Gusic, Zeljko Kucan, Carles Lalueza-Fox, Marco de la Rasilla, Antonio Rosas, Pavao Rudan, Sandra Knapp, Bruce Benz, Michael Blake, R.G. Matson, Bryn Dentinger, Anna Stalter, Robert Capers, John Peter for providing samples.

---

**Figure S1.**
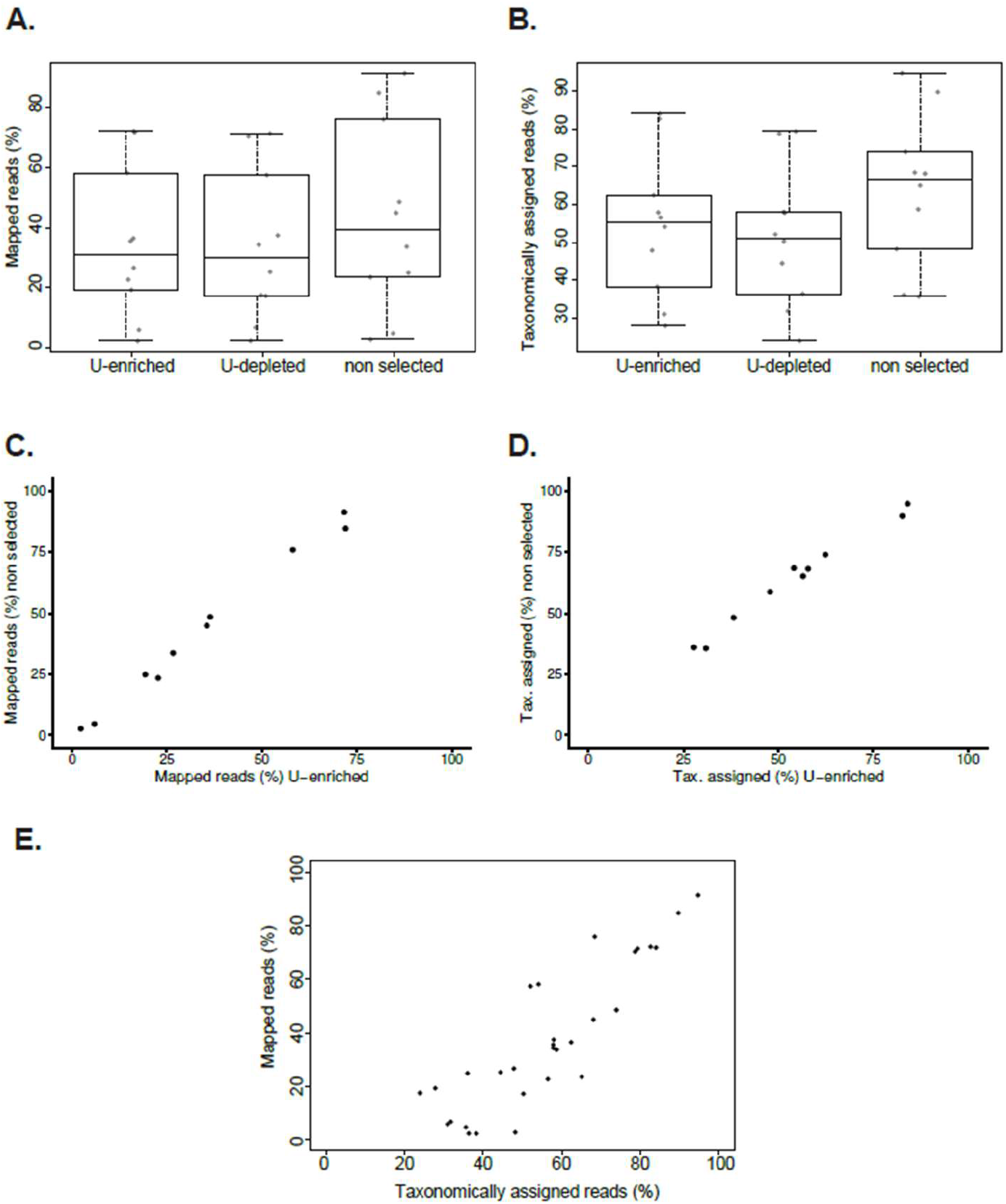
Mapped and taxonomically assigned reads of plant historical specimens. **A.** Distributions of percentage of mapped reads for non-selected and U-selected libraries (U-enriched and U-depleted fractions). **B.** Distributions of percentage of taxonomically assigned reads for nonselected and U-selected libraries (U-enriched and U-depleted fractions). **C.** Correlation of the percentage of mapped reads between the U-enriched and the non-selected library **D.** Correlation of the percentage of taxonomically assigned reads between the U-enriched and the non-selected library **E.** Relation between percentages of mapped and taxonomically assigned reads from U-selected libraries (U-enriched fraction).

**Figure S2.**
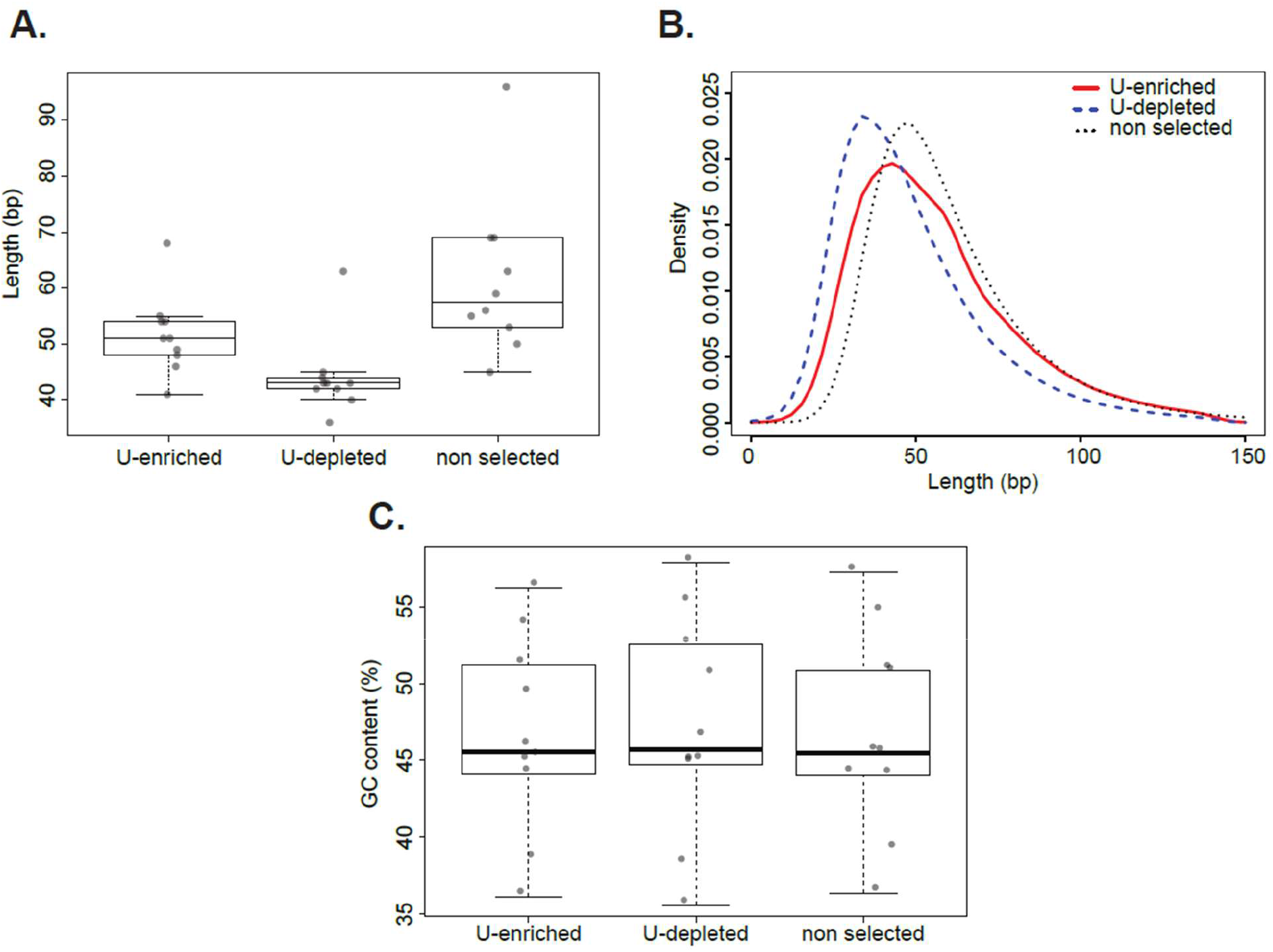
Length and GC content of plant historical specimens. **A.** Distributions of mean length for non-selected and U-selected libraries (U-enriched and U-depleted fractions). Median values are denoted as black lines and points show the original value for each individual sample. **B.** Length distribution of *Arabidopsis thaliana* sample NY1365375 for a non-selected and U-selected library (U-enriched and U-depleted fractions). **C.** Distributions of mean GC content for non-selected and U-selected libraries (U-enriched and U-depleted fractions).

**Figure S3.**
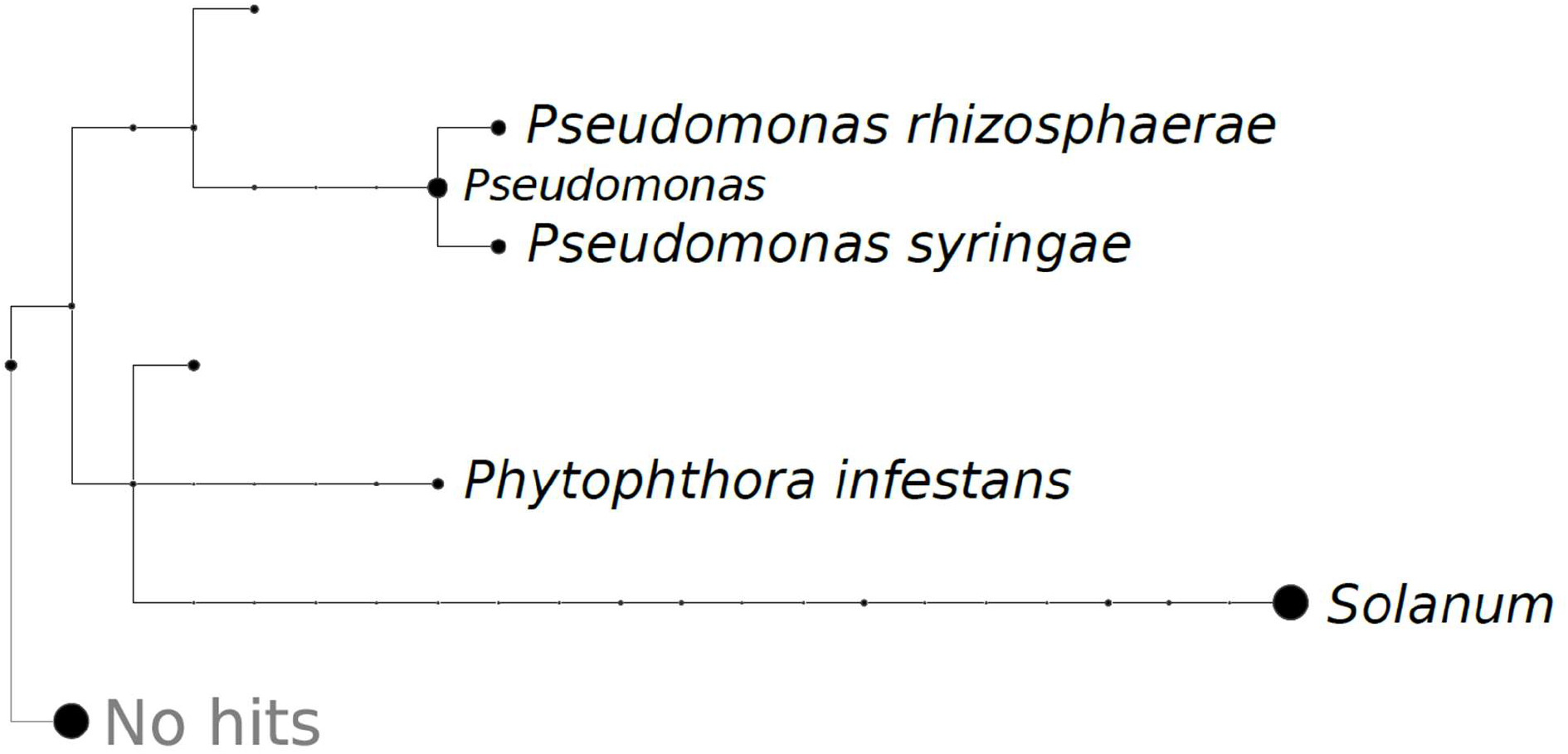
Taxonomic tree of reads from a *Solanum tuberosum* library assigned to different taxonomic levels. The size of the circle represents the amount of reads assigned to the node displayed in the tree or to any taxonomic level below it. Reads assigned to some species *Phytophthora infestans, Pseudomonas syringae and Pseudomonas rhizosphaerae*, as well as the genera Pseudomonas and Solanum are named in the taxonomic tree.

**Figure S4.**
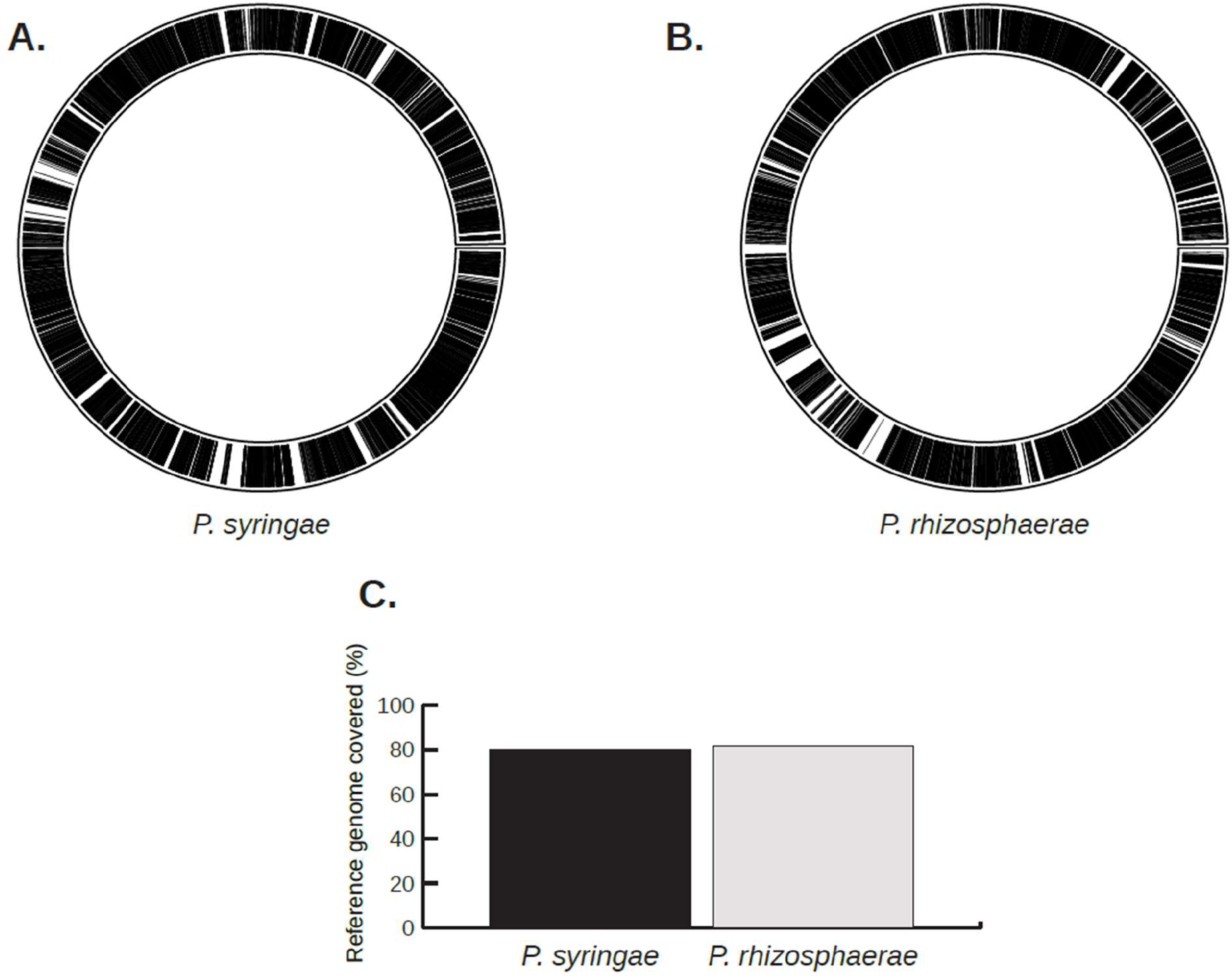
Analysis of de novo assembled *Pseudomonas* contigs from a *Solanum tuberosum* sample. Contig coverage of *Pseudomonas syringae* (**A.**) and *Pseudomonas rhizosphaerae* (**B.**). The circles represent each circular bacterial reference genome, and the black lines depict genomic regions covered by alignments of de novo assembled contigs. **C.** Percentage of reference genome of *P. syringae* and *P. rhizosphaerae* covered by de novo assembled contigs from A. and B., respectively.

**Figure S5.**
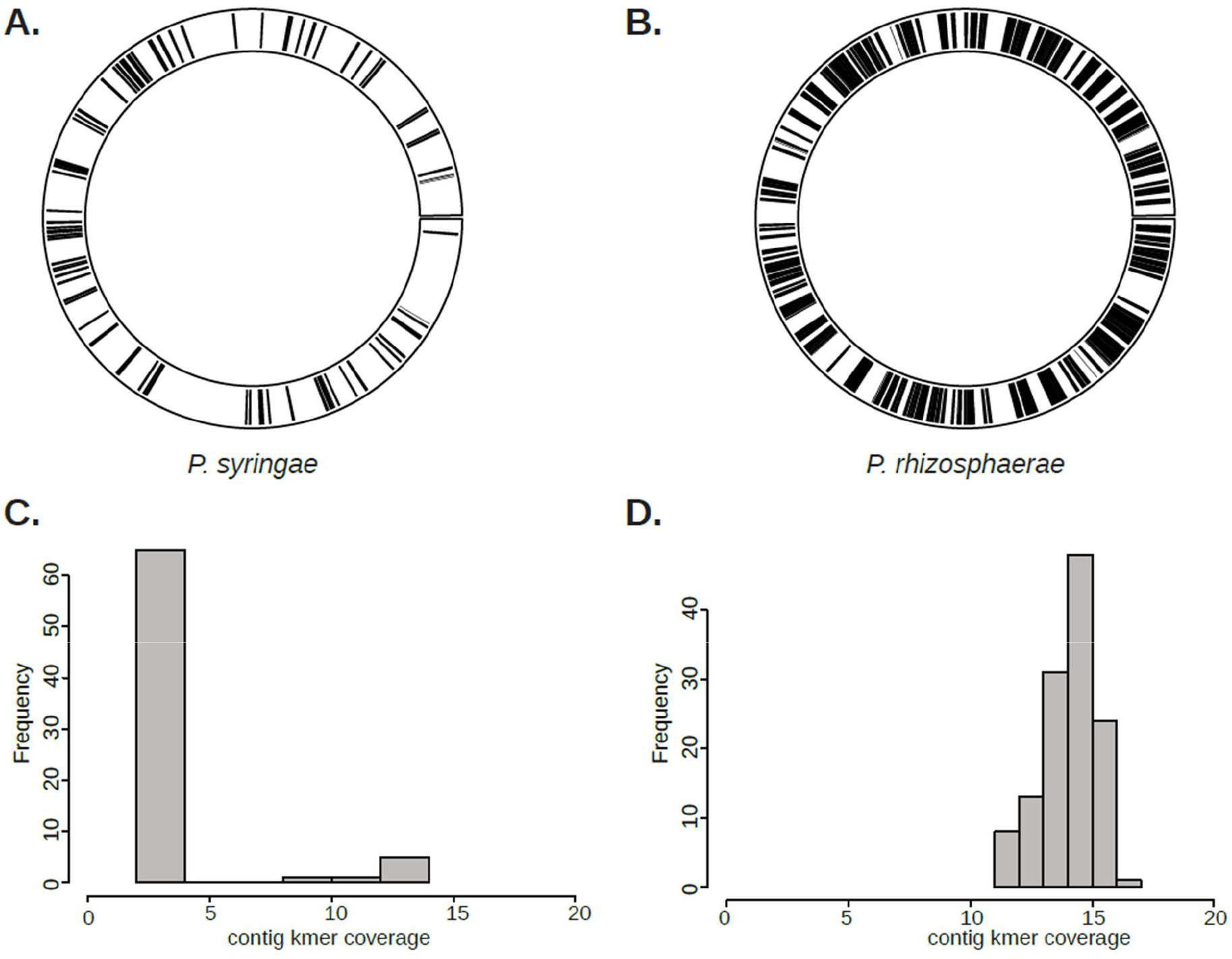
Analysis of uniquely mapped *Pseudomonas* contigs from a *Solanum tuberosum* sample. Uniquely mapped contig coverage of *Pseudomonas syringae* (**A.**) and *Pseudomonas rhizosphaerae* (**B.**). The circles represent each circular bacterial reference genome, and the black lines depict genomic regions covered by alignments of uniquely mapped contigs. Histograms of contig k-mer coverage from de novo assembled contigs uniquely mapping to *P. syringae* (**C.**) and *P. rhizosphaerae* (**D.**).

## References

1. Poinar HN, Schwarz C, Qi J, Shapiro B, Macphee RDE, Buigues B, et al. Metagenomics to paleogenomics: large-scale sequencing of mammoth DNA. Science. 2006;311:392–4. doi:10.1126/science.1123360.

2. Gutaker RM, Burbano HA. Reinforcing plant evolutionary genomics using ancient DNA. Curr Opin Plant Biol. 2017;36:38–45. doi:10.1016/j.pbi.2017.01.002.

3. Orlando L, Gilbert MTP, Willerslev E. Reconstructing ancient genomes and epigenomes. Nat Rev Genet. 2015;16:395–408. doi:10.1038/nrg3935.

4. Warinner C, Herbig A, Mann A, Fellows Yates JA, Weiß CL, Burbano HA, et al. A Robust Framework for Microbial Archaeology. Annu Rev Genomics Hum Genet. 2017;18:321–56. doi:10.1146/annurev-genom-091416-035526.

5. Bos KI, Schuenemann VJ, Golding GB, Burbano HA, Waglechner N, Coombes BK, et al. A draft genome of Yersinia pestis from victims of the Black Death. Nature. 2011;478:506–10. doi:10.1038/nature10549.

6. Yoshida K, Schuenemann VJ, Cano LM, Pais M, Mishra B, Sharma R, et al. The rise and fall of the Phytophthora infestans lineage that triggered the Irish potato famine. Elife. 2013;2:e00731. doi:10.7554/eLife.00731.

7. Adler CJ, Dobney K, Weyrich LS, Kaidonis J, Walker AW, Haak W, et al. Sequencing ancient calcified dental plaque shows changes in oral microbiota with dietary shifts of the Neolithic and Industrial revolutions. Nat Genet. 2013;45:450-5, 455e1. doi:10.1038/ng.2536.

8. Warinner C, Rodrigues JFM, Vyas R, Trachsel C, Shved N, Grossmann J, et al. Pathogens and host immunity in the ancient human oral cavity. Nat Genet. 2014;46:336–44. doi: 10.1038/ng.2906.

9. Weyrich LS, Duchene S, Soubrier J, Arriola L, Llamas B, Breen J, et al. Neanderthal behaviour, diet, and disease inferred from ancient DNA in dental calculus. Nature. 2017. doi:10.1038/nature21674.

10. Boast AP, Weyrich LS, Wood JR, Metcalf JL, Knight R, Cooper A. Coprolites reveal ecological interactions lost with the extinction of New Zealand birds. Proc Natl Acad Sci U S A. 2018;115:1546–51. doi:10.1073/pnas.1712337115.

11. Briggs AW, Stenzel U, Johnson PLF, Green RE, Kelso J, Prüfer K, et al. Patterns of damage in genomic DNA sequences from a Neandertal. Proc Natl Acad Sci U S A. 2007; 104:14616–21. doi:10.1073/pnas.0704665104.

12. Krause J, Briggs AW, Kircher M, Maricic T, Zwyns N, Derevianko A, et al. A complete mtDNA genome of an early modern human from Kostenki, Russia. Curr Biol. 2010;20:231–6. doi:10.1016/j.cub.2009.11.068.

13. Prüfer K, Meyer M. Anthropology. Comment on “Late Pleistocene human skeleton and mtDNA link Paleoamericans and modern Native Americans.” Science. 2015;347:835. doi:10.1126/science.1260617.

14. Weiß CL, Dannemann M, Prüfer K, Burbano HA. Contesting the presence of wheat in the British Isles 8,000 years ago by assessing ancient DNA authenticity from low-coverage data. Elife. 2015;4. doi:10.7554/eLife.10005.

15. Gansauge M-T, Meyer M. Selective enrichment of damaged DNA molecules for ancient genome sequencing. Genome Res. 2014;24:1543–9. doi:10.1101/gr.174201.114.

16. Skoglund P, Malmström H, Raghavan M, Storå J, Hall P, Willerslev E, et al. Origins and genetic legacy of Neolithic farmers and hunter-gatherers in Europe. Science. 2012;336:466–9. doi:10.1126/science.1216304.

17. Meyer M, Fu Q, Aximu-Petri A, Glocke I, Nickel B, Arsuaga J-L, et al. A mitochondrial genome sequence of a hominin from Sima de los Huesos. Nature. 2014;505:403–6. doi: 10.1038/nature12788.

18. Zaremba-Niedźwiedzka K, Andersson SGE. No ancient DNA damage in Actinobacteria from the Neanderthal bone. PLoS One. 2013;8:e62799. doi:10.1371/journal.pone.0062799.

19. Weiß CL, Schuenemann VJ, Devos J, Shirsekar G, Reiter E, Gould BA, et al. Temporal patterns of damage and decay kinetics of DNA retrieved from plant herbarium specimens. R Soc Open Sci. 2016;3:160239. doi:10.1098/rsos.160239.

20. Meyer M, Kircher M. Illumina sequencing library preparation for highly multiplexed target capture and sequencing. Cold Spring Harb Protoc. 2010;2010:db.prot5448. doi:10.1101/pdb.prot5448.

21. Gansauge M-T, Meyer M. Single-stranded DNA library preparation for the sequencing of ancient or damaged DNA. Nat Protoc. 2013;8:737–48. doi:10.1038/nprot.2013.038.

22. Gansauge M-T, Gerber T, Glocke I, Korlevic P, Lippik L, Nagel S, et al. Singlestranded DNA library preparation from highly degraded DNA using T4 DNA ligase. Nucleic Acids Res. 2017;45:e79. doi:10.1093/nar/gkx033.

23. Smits THM, Rezzonico F, Kamber T, Goesmann A, Ishimaru CA, Stockwell VO, et al. Genome sequence of the biocontrol agent Pantoea vagans strain C9-1. J Bacteriol. 2010;192:6486–7. doi:10.1128/JB.01122-10.

24. Exposito-Alonso M, Becker C, Schuenemann VJ, Reiter E, Setzer C, Slovak R, et al. The rate and potential relevance of new mutations in a colonizing plant lineage. PLoS Genet. 2018;14:e1007155. doi:10.1371/journal.pgen.1007155.

25. Swarts K, Gutaker RM, Benz B, Blake M, Bukowski R, Holland J, et al. Genomic estimation of complex traits reveals ancient maize adaptation to temperate North America. Science. 2017;357:512–5. doi:10.1126/science.aam9425.

26. Kircher M, Sawyer S, Meyer M. Double indexing overcomes inaccuracies in multiplex sequencing on the Illumina platform. Nucleic Acids Res. 2012;40:e3. doi:10.1093/nar/gkr771.

27. Briggs AW, Stenzel U, Meyer M, Krause J, Kircher M, Pääbo S. Removal of deaminated cytosines and detection of in vivo methylation in ancient DNA. Nucleic Acids Res. 2010;38:e87. doi:10.1093/nar/gkp1163.

28. Kircher M. Analysis of high-throughput ancient DNA sequencing data. Methods Mol Biol. 2012;840:197–228. doi:10.1007/978-1-61779-516-9_23.

29. Renaud G, Stenzel U, Kelso J. leeHom: adaptor trimming and merging for Illumina sequencing reads. Nucleic Acids Res. 2014;42:e141. doi:10.1093/nar/gku699.

30. Schnable PS, Ware D, Fulton RS, Stein JC, Wei F, Pasternak S, et al. The B73 maize genome: complexity, diversity, and dynamics. Science. 2009;326:1112–5. doi:10.1126/science.1178534.

31. Arabidopsis Genome Initiative. Analysis of the genome sequence of the flowering plant Arabidopsis thaliana. Nature. 2000;408:796–815. doi:10.1038/35048692.

32. Swarbreck D, Wilks C, Lamesch P, Berardini TZ, Garcia-Hernandez M, Foerster H, et al. The Arabidopsis Information Resource (TAIR): gene structure and function annotation. Nucleic Acids Res. 2008;36 Database issue:D1009-14. doi:10.1093/nar/gkm965.

33. Potato Genome Sequencing Consortium, Xu X, Pan S, Cheng S, Zhang B, Mu D, et al. Genome sequence and analysis of the tuber crop potato. Nature. 2011;475:189–95. doi:10.1038/nature10158.

34. Tomato Genome Consortium. The tomato genome sequence provides insights into fleshy fruit evolution. Nature. 2012;485:635–41. doi:10.1038/nature11119.

35. Li H. Aligning sequence reads, clone sequences and assembly contigs with BWA-MEM. arXiv [q-bio.GN]. 2013. http://arxiv.org/abs/1303.3997.

36. Herbig A, Maixner F, Bos KI, Zink A, Krause J, Huson DH. MALT: Fast alignment and analysis of metagenomic DNA sequence data applied to the Tyrolean Iceman. bioRxiv. 2016;:050559. doi:10.1101/050559.

37. Huson DH, Auch AF, Qi J, Schuster SC. MEGAN analysis of metagenomic data. Genome Res. 2007;17:377–86. doi:10.1101/gr.5969107.

38. Nolan M, Sikorski J, Jando M, Lucas S, Lapidus A, Del Rio TG, et al. Complete genome sequence of Streptosporangium roseum type strain (NI 9100 T). Stand Genomic Sci. 2010;2:29. https://standardsingenomics.biomedcentral.com/articles/10.4056/sigs.631049.

39. Feil H, Feil WS, Chain P, Larimer F, DiBartolo G, Copeland A, et al. Comparison of the complete genome sequences of Pseudomonas syringae pv. syringae B728a and pv. tomato DC3000. Proc Natl Acad Sci U S A. 2005;102:11064–9. doi:10.1073/pnas.0504930102.

40. Kwak Y, Jung BK, Shin J-H. Complete genome sequence of Pseudomonas rhizosphaerae IH5 T (= DSM 16299 T), a phosphate-solubilizing rhizobacterium for bacterial biofertilizer. J Biotechnol. 2015;193:137–8. http://www.sciencedirect.com/science/article/pii/S0168165614010293.

41. Langmead B, Salzberg SL. Fast gapped-read alignment with Bowtie 2. Nat Methods. 2012;9:357–9. doi: 10.1038/nmeth.1923.

42. Jónsson H, Ginolhac A, Schubert M, Johnson PLF, Orlando L. mapDamage2.0: fast approximate Bayesian estimates of ancient DNA damage parameters. Bioinformatics. 2013;29:1682–4. doi: 10.1093/bioinformatics/btt193.

43. Li H. A statistical framework for SNP calling, mutation discovery, association mapping and population genetical parameter estimation from sequencing data. Bioinformatics. 2011;27:2987–93. doi:10.1093/bioinformatics/btr509.

44. Li H. Minimap2: pairwise alignment for nucleotide sequences. Bioinformatics. 2018;34:3094–100. doi:10.1093/bioinformatics/bty191.

45. Knaus BJ, Grünwald NJ. vcfr: a package to manipulate and visualize variant call format data in R. Mol Ecol Resour. 2017;17:44–53. doi:10.1111/1755-0998.12549.

46. Jombart T, Ahmed I. adegenet 1.3-1: new tools for the analysis of genome-wide SNP data. Bioinformatics. 2011;27:3070–1. doi:10.1093/bioinformatics/btr521.

47. R Development Core Team. R: A Language and Environment for Statistical Computing. 2008. http://www.R-project.org.

48. Bankevich A, Nurk S, Antipov D, Gurevich AA, Dvorkin M, Kulikov AS, et al. SPAdes: a new genome assembly algorithm and its applications to single-cell sequencing. J Comput Biol. 2012;19:455–77. doi:10.1089/cmb.2012.0021.

49. Harris RS. Improved pairwise alignment of genomic DNA. The Pennsylvania State University; 2007. http://search.proquest.com/openview/bc77cca0fb9390b44b9ef572fb574322/1?pq-origsite=gscholar&cbl=18750&diss=y.

50. Krzywinski M, Schein J, Birol I, Connors J, Gascoyne R, Horsman D, et al. Circos: an information aesthetic for comparative genomics. Genome Res. 2009;19:1639–45. doi:10.1101/gr.092759.109.

51. Hohlfeld S, Ankenbrand M, Förster F, Hackl T. AliTV: Version 0.4.1. Zenodo; 2016. doi:10.5281/zenodo.59682.

52. Gu Z, Gu L, Eils R, Schlesner M, Brors B. circlize Implements and enhances circular visualization in R. Bioinformatics. 2014;30:2811–2. doi:10.1093/bioinformatics/btu393.

